# Diel CO_2_ fluctuations alter the molecular response of coral reef fishes to ocean acidification conditions

**DOI:** 10.1101/2021.05.17.444406

**Authors:** Celia Schunter, Michael D. Jarrold, Philip L. Munday, Timothy Ravasi

## Abstract

Environmental CO_2_ variation can modify the responses of marine organisms to ocean acidification, yet the underlying mechanisms for this effect remain unclear. On coral reefs, environmental CO_2_ fluctuates on a regular day-night cycle. Effects of future ocean acidification on coral reef fishes might therefore depend on their response to this diel cycle of CO_2_. To evaluate the effects on the brain molecular response, we exposed two common reef fishes (*Acanthochromis polyacanthus* and *Amphiprion percula*) to two projected future CO_2_ levels (750 and 1,000 μatm) under both stable and diel fluctuating conditions. We found a common signature to stable elevated pCO_2_ for both species, which included the downregulation of immediate early genes, indicating lower brain activity. The transcriptional program was more strongly affected by higher average CO_2_ in a stable treatment than for fluctuating treatments, however, the largest difference in molecular response was between stable and fluctuating CO_2_ treatments. This indicates that a response to a change in environmental CO_2_ conditions is different for organisms living in a fluctuating than in stable environments. The differential regulation was related to steroid hormones and circadian rhythm (CR). Both species exhibited a marked difference in the expression of CR genes among CO_2_ treatments, possibly accommodating a more flexible adaptive approach to acid-base control, which could explain reduced impairment. Our results suggest that environmental CO_2_ fluctuations might enable reef fishes to phase shift their clocks and anticipate CO_2_ changes, thereby avoiding impairments and more successfully adjust to ocean acidification conditions.

## Introduction

How organisms respond to natural environmental variation can affect their ability to cope with human-induced environmental change (Lawson, Vindenes, Bailey, & van de Pol, 2015). The physiological, behavioural and molecular responses of marine organisms to predicted future climate change have been intensely investigated in the last decade (Fuller et al., 2010; Strader, Wong, & Hofmann, 2020; Wong & Candolin, 2015), yet the role of natural environmental variation is often overlooked (Brown et al., 2016; Vajedsamiei, Wahl, Schmidt, Yazdanpanahan, & Pansch, 2021). Organisms are adapted to the environmental variation they experience in their natural habitats (Melbinger & Vergassola, 2015), which in turn can influence their physiological responses and adaptive capacity to environmental change (Kroeker et al., 2020). Yet, natural variability in environmental conditions has often been ignored when experimentally evaluating species responses to global change, with a primary focus on how the average conditions may change, without also considering how the variation in those conditions may change. Organisms have evolved buffering mechanisms, including behaviour or physiological adjustments, to deal with natural environmental fluctuations (Bernhardt, O’Connor, Sunday, & Gonzalez, 2021; Huey & Bennett, 1990) and for marine fishes physiological responses and behaviour, among others, have shown to vary depending on fluctuations in environmental conditions (Claireaux, Webber, Kerr, & Boutilier, 1995; Enders & Boisclair, 2016; Huey & Bennett, 1990; Villegas-Ríos, Réale, Freitas, Moland, & Olsen, 2018). Despite the importance of environmental variability to organisms’ responses to climate change there is a lack of information on the molecular mechanisms driving these adjustments. Importantly, in the light of predicted increasing environmental variability associated with climate change, we need to incorporate such variation into experiments to better evaluate the ecological consequences of climate change in both marine and terrestrial ecosystems (Rivest, Comeau, & Cornwall, 2017; Stenseth et al., 2002).

One aspect of global change is the increased oceanic uptake of carbon dioxide (CO_2_) from the atmosphere causing the partial pressure of CO_2_ (pCO_2_) at the ocean surface to increase and the pH of seawater to decline, a process called ocean acidification (OA; Doney, Fabry, Feely, & Kleypas, 2009). Ocean acidification projections are based on data from open ocean environments that have relatively stable *p*CO_2_ through time (Doney, Busch, Cooley, & Kroeker, 2020; Hofmann et al., 2011). However, when open ocean waters flush onto shallow water coastal habitats their carbonate chemistry can be substantially modified by a range of biological and physical processes, resulting in significant fluctuations in *p*CO_2_ on a variety of temporal scales (Duarte, Losada, Hendriks, Mazarrasa, & Marbà, 2013; Hendriks et al., 2015; Hofmann et al., 2011; Waldbusser & Salisbury, 2014). Continuous time series data has revealed that *p*CO_2_ fluctuations in such habitats often exceed those in the open ocean by an order of magnitude (Hofmann et al., 2011; Shaw, McNeil, & Tilbrook, 2012). In some instances, these fluctuations can even exceed the mean CO_2_ levels projected to occur over the next century (Baumann, Wallace, Tagliaferri, & Gobler, 2015; Duarte et al., 2013; Shaw et al., 2012). Furthermore, the magnitude of natural *p*CO_2_ fluctuations is expected to increase throughout the century, as increased CO_2_ uptake by the oceans leads to reduced seawater buffering capacity (Pacella, Brown, Waldbusser, Labiosa, & Hales, 2018; Schulz & Riebesell, 2013; Shaw, McNeil, Tilbrook, Matear, & Bates, 2013). Yet, how natural *p*CO_2_ fluctuations affect the biological responses of marine organism to higher average CO_2_ levels in the ocean is still relatively unknown.

A large body of experimental research has shown that OA poses a threat to shallow water marine organisms, impacting a range of biological traits (Cattano, Claudet, Domenici, & Milazzo, 2018; Kroeker et al., 2013; Przeslawski, Byrne, & Mellin, 2015; Wittmann & Pörtner, 2013). However, most laboratory experiments on shallow water marine organisms have been conducted using stable levels of elevated *p*CO_2_ consistent with open ocean projections, instead of those ecological relevant to the study organism (Gunderson, Armstrong, & Stillman, 2016; McElhany & Shallin Busch, 2013; Wahl, Saderne, & Sawall, 2016). Empirical studies testing populations across their geographic range, have shown that sensitivity to future stable ocean acidification conditions is linked to the local *p*CO_2_ conditions experienced (Hoshijima & Hofmann, 2019; Kelly & Hofmann, 2013; Pespeni et al., 2013; Thomsen et al., 2017; Vargas et al., 2017). Perhaps the most well-known natural *p*CO_2_ fluctuations in shallow water coastal habitats, including coral reefs, macroalgal beds, seagrass meadows and salt marshes are those that occur across a 24-hour period (Baumann et al., 2015; Challener, Robbins, & McClintock, 2016; Kline et al., 2015b; Wahl et al., 2018). These daily CO_2_ cycles are driven by the processes of photosynthesis/respiration and calcification/dissolution over a day-night cycle, but are modified by physical properties such as flow trajectory, flow rates and residence time, which ultimately alters the contact time between the seawater and the benthic community (Cyronak et al., 2018; Enochs et al., 2020, 2018; Falter, Lowe, Zhang, & McCulloch, 2013; Shaw et al., 2012). Importantly, a number of more recent laboratory studies have shown that natural diel CO_2_ cycles can significantly modify the biological responses (e.g. growth, survival, behaviour and metabolism) of shallow water marine organisms to OA (Cornwall et al., 2013; Enochs et al., 2018; Frieder, Gonzalez, Bockmon, Navarro, & Levin, 2014; Jarrold, Humphrey, McCormick, & Munday, 2017; Mangan, Urbina, Findlay, Wilson, & Lewis, 2017; Ou et al., 2015; Wahl et al., 2018). Establishing the underlying mechanisms responsible for the effects CO_2_ variability has on OA sensitivity will be critical for predicting the effects of future ocean acidification in natural marine habitats where environmental CO_2_ fluctuates on a range on spatial and temporal scales.

Marine fish were initially expected to be resilient to OA as they can tightly defend their internal pH under elevated CO_2_ conditions *via* active control of acid-base relevant ions (Baker et al., 2009; R. M. Heuer & Grosell, 2014; Wood, Turner, Munger, & Graham, 1990). However, experimental studies have reported significant effects of elevated CO_2_ on growth, survival, physiological condition, otolith calcification and behaviour, especially in early life stages (Baumann, 2019; Cattano et al., 2018; Munday, Jarrold, & Nagelkerken, 2019). In particular, over 80 studies, on a range of tropical and temperate species, have shown that exposure to stable *p*CO_2_ levels above 700 μatm can affect cognitive processes, impair a range of sensory systems and alter ecologically important behaviours (Cattano et al., 2018; Munday et al., 2020, 2019; Paula, Repolho, et al., 2019). These impairments are thought to be linked to the process of regulating acid-base balance in a high CO_2_ environment (Rachael M. Heuer, Hamilton, & Nilsson, 2019).

Fish adjust the concentrations of acid-base relevant ions (primarily HCO_3_ and Cl^-^) to maintain homeostasis of internal pH in high CO_2_ conditions (Esbaugh, 2018) including in the brain (Esbaugh, Ern, Nordi, & Johnson, 2016; Rachael M. Heuer & Grosell, 2014). These ionic changes could affect the function of GABA-A receptors in the brain, causing behavioural changes (Nilsson et al., 2012; Schunter, Ravasi, Munday, & Nilsson, 2019). However, not all species are equally affected and numerous studies have reported no effects of OA conditions on fish behaviours (Clark et al., 2020; Heinrich et al., 2015; Jutfelt & Hedgärde, 2013; Sundin, Vossen, Nilsson-Sköld, & Jutfelt, 2017). While species and/or trait specific response and differences in methods used may explain contrasting results between studies (Munday et al., 2020, 2019), it is also possible that different results might partly be explained by differences in the stability of *p*CO_2_ treatments in laboratory experiments. Indeed, recent studies have shown that diel fluctuations in CO_2_ can mitigate and alleviate behavioural impairments caused by stable elevated CO_2_ (Jarrold et al., 2017; Laubenstein, Jarrold, Rummer, & Munday, 2020). Fitness related traits such as larval survival and growth, as well as energy consumption, also improved with diel CO_2_ fluctuations, suggesting with these environmental fluctuations might provide a temporary physiological refuge from persistent high pCO_2_ (Cross, Murray, & Baumann, 2019; Hannan, Munday, & Rummer, 2020; Jarrold & Munday, 2019; Ou et al., 2015). Yet how these adjustments to diel changes in pCO_2_ are regulated, and how fish adjust to environmental fluctuations in pCO_2_ on the molecular level it currently unknown. The response of coral reef organisms to future ocean acidification may be contingent on their physiological responses to natural environmental CO_2_ variation, and may not be well described by experiments using stable elevated CO_2_ treatments.

With environmental fluctuation predicted to increase as climate change progresses (McNeil & Sasse, 2016; Vázquez, Gianoli, Morris, & Bozinovic, 2017) it is essential to understand the underlying mechanisms that marine organisms use to deal with environmental variability. Jarrold et al. (2017) showed that diel CO_2_ cycles could either partially or fully alleviated the negative effects of OA on behavioural impairments in two common coral reef fishes, *Acanthochromis polyacanthus* and *Amphiprion percula*, dependent on the mean CO_2_ and magnitude of fluctuation experienced (Supplementary Figure S1). Here we tested the effects of stable and fluctuating CO_2_ treatments on the transcriptional response in the brain of the fish in that experiment. Both species were reared in three stable (460, 750 and 1,000 μatm) and three fluctuating (550 ± 150, 750 ± 300 and 1,000 ± 300 μatm) CO_2_ treatments (Figure 1). The brain transcriptomes where then compared among treatments in order to determine how molecular processes differ between fish in stable versus fluctuating high CO_2_ environments. Thus, here, we examined the transcriptional brain response of fishes from the Jarrold et al. (2017) experiments to test the effects of diel CO_2_ fluctuations on the molecular brain responses of coral reef fishes to different ocean acidification scenarios.

**Fig. 1:**
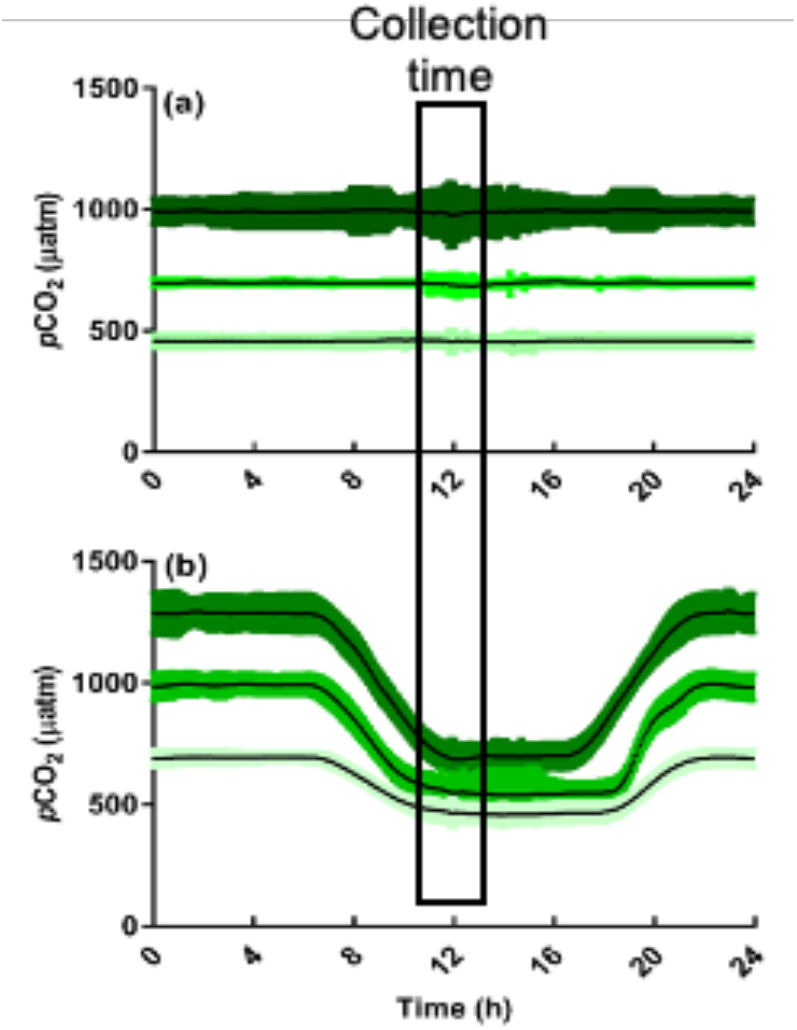
Mean diel stable (above) and fluctuating (below) pCO_2_ profiles. Coloured sections are ± 1 SD. The black window illustrates when fish were sampled for brain transcriptome analysis.

The stable 1,000 μatm *p*CO_2_ treatment represented the open ocean projection for the end of this century typically used in many OA experiments (Kroeker et al., 2013). The magnitude of variation in the fluctuating CO_2_ treatments (± 300 μatm) matched levels that have been observed in some tidal lagoons (Shaw et al., 2012). Diel *p*CO_2_ fluctuations of between ± 50-150 μatm are more typical in most other reef areas where the species used in this study reside (Albright, Langdon, & Anthony, 2013; Hannan, Miller, et al., 2020; Kline et al., 2015a). However, the magnitude of CO_2_ fluctuations is predicted to increase at least three-fold by the year 2100 as OA leads to the reduced buffering capacity of seawater (McNeil & Matsumoto, 2019; McNeil & Sasse, 2016; Shaw et al., 2013) and thus the species used in this study could be exposed to daily fluctuations of between 150-450 μatm by the end of the century. Previous experiments indicate that behavioural abnormalities are first evident at around 700 μatm CO_2_, although the magnitude of effect is not as large as observed at higher CO_2_ levels (Ferrari et al., 2011; Jarrold et al., 2017; Munday et al., 2010b; Paula, Baptista, et al., 2019; Welch, Watson, Welsh, McCormick, & Munday, 2014). Therefore, the inclusion of the 750 and 750 ± 300 μatm CO_2_ treatments enabled us to determine how diel *p*CO_2_ cycles may affect the onset threshold of behavioural abnormalities and the molecular responses in the brains of fishes. With environmental fluctuation predicted to increase as climate change progresses (McNeil & Sasse, 2016; Vázquez et al., 2017) it is essential to understand the underlying mechanisms that marine organisms use to deal with environmental variability.

## Materials and Methods

### Brood-stock and general rearing protocol

The two species of damselfish (family Pomacentridae) in this study, *Acanthochromis polyacanthus* and *Amphiprion percula*, are commonly found on coral reefs in the Great Barrier Reef. After hatching from benthic eggs, *Am. percula* has a pelagic larval phase of approximately 11 days, followed by benthic juvenile and adult stages. By contrast, *Ac. polyacanthus* does not have a pelagic larval stage and instead has direct developing juveniles that remain in the reef habitat from hatching. Therefore, these two species could potentially have different adaptations to environmental CO_2_ variation in their early life stages.

Adult *Ac. polyacanthus* were collected using hand nets from the Bramble Reef area (site 1: 18°22’S, 146°40’E; site 2: 18°25’S, 146°40’E) of the Great Barrier Reef in July 2015. Fish were transported to an environmentally controlled aquarium research facility at James Cook University (JCU; Townsville, Australia) where they were housed as breeding pairs in 60 L aquaria at temperature conditions matching the collection location. An existing brood-stock of wild-caught (in 2017) *Am. percula* at JCU was used. These fish had been collected from the Cairns Region of the Great Barrier Reef and housed at JCU for four years. Therefore, while collected at different times and locations on the Great Barrier Reef, the breeding pairs for both species were wild caught fish. Adult *Ac. polyacanthus* and *Am. percula* pairs were maintained under stable ambient *p*CO_2_ (~490 μatm). Water temperature was increased at a rate of 0.5°C *per* week until the summer breeding temperature of 29°C was reached in the first week of November 2015. Adult pairs were provided with half a terracotta pot to act as a shelter and spawning site. Aquaria were checked each morning for the presence of newly laid clutches. Pairs were fed *ad libitum* on commercial fish feed pellets (INVE Aquaculture Nutrition NRD 12/20) once daily outside the breeding season and twice daily during the breeding season (November-May). *Ac. polyacanthus* juveniles were fed a combination of freshly hatched *Artemia* naupli and weaning fish feed (INVE Aquaculture Nutrition Wean-S 200-400 μm) for the first four days post hatch (dph).

From 5-21 dph they were fed daily on the weaning feed and then switched to a small pellet fish feed (INVE Aquaculture Nutrition NRD 5/8) at 22 dph. Rearing of larval *Am. percula* was performed using methods described by Jarrold et al. (2017). Postsettlement stage juveniles were fed daily on the weaning fish feed until 21 dph and were then switched to a small pellet fish feed at 22 dph.

### Experimental design

The experiment was carried out at the National Sea Simulator (SeaSim) facility at the Australian Institute of Marine Science (AIMS) (Cape Cleveland, Australia). Offspring (i.e. F1 generation) from three clutches of *Ac. polyacanthus* and three clutches of *Am. percula* were transferred to the experimental system and each clutch was split between all the *p*CO_2_ treatments in duplicate tanks per treatment. Each clutch was from a different parental pair. Each tank was stocked with 15 fish for *Ac. polyacanthus* and 13-15 fish for *Am. percula* at 14 and 12 dph respectively. Fish were reared in experimental treatments until the age of 10 weeks. Behavioural tests including lateralization and the response to chemical alarm cues performed as part of this experiment were previously reported by Jarrold et al. (2017; Supplementary Figure S1).

### Experimental system and CO_2_ manipulation

The experimental system used at SeaSim was a flow-through system which comprised of multiple independent lines (duplicate independent lines per *p*CO_2_ treatment). The system used ultra-filtered seawater (0.04 μm); temperature controlled to 28.5°C. Each seawater line supplied three custom made 50 L tanks at the rate of 50 L h^-1^. The experimental tanks were placed in individual temperature-controlled water baths to ensure temperature stability (± 0.1°C). Treatments and tank replicates were randomly positioned in the experimental room. The management of *p*CO_2_ and temperature was achieved through the design and implementation of a custom Model Predictive Control logic running on a micro-PLC (Series S7-1500, Siemens, Australia). The micro-PLC was integrated with the general SeaSim control system, to provided SCADA (Siemens WinCC) accessibility and data archiving. The *p*CO_2_ feedback for each of the replication lines was provided via non-dispersive infrared measurements (Dixson, Sabine, & Christian, 2007). Tank water was delivered to the equilibrator (SeaSim, AIMS design, custom built) by an in-tank submersible pump (Universal Pump 1260, EHEIM, Deizisau, Germany) where the *p*CO_2_ of the air in the chamber reaches and maintains equilibrium with the *p*CO_2_ of the experimental water. The air was constantly delivered to a NDIR CO_2_ analyser (T elaire T6613, Amphenol, Australia) that provided live feedback to the PLC. The CO_2_ analysers were calibrated monthly using certified calibration gas mixtures at 0, 600 and 2000 ppm. The control system delivered CO_2_ though Gas Mass Flow Controllers (GFC17 series, Aalborg, Orangeburg, USA) according to the profiling schedule designed for the *p*CO_2_ treatment and the feedback signal coming from the experimental tanks. CO_2_ was dissolved in the flow-through water by mean of membrane contactors (Membrana Liqui-Cel 2.5×8 Extra-Flow. 3M, USA). Total alkalinity was also measured weekly as described above.

The magnitude and durations of diel CO_2_ fluctuations on coral reefs vary spatially and temporally due to physical forcing from wind and tidal currents (Cyronak et al., 2018; Falter et al., 2013; Hannan, Miller, et al., 2020), and are more extreme in lagoonal areas than offshore reefs flushed by oceanic waters (Albright et al., 2013; Shaw et al., 2013). We used a diel CO_2_ cycle based on that observed on coral reefs in the Great Barrier Reef (Albright et al., 2013; Hannan, Miller, et al., 2020). For logistical reasons the duration of our high and low CO_2_ peaks were longer that observed on natural coral reefs, but the times at which CO_2_ began to fall and rise after each peak were similar to that observed in nature (Figure 1).

Throughout the experiment incoming coastal water had a *p*CO_2_ ranging between 500-550 μatm. Thus, to achieve a control *p*CO_2_ level closer to atmospheric levels (~400 μatm, membrane contactors (Membrana Liqui-Cel 4×28 Extra-Flow) were used to remove CO_2_, using CO_2_-depleted air as sweep gas. Using this method, we were able to achieve a stable control treatment with a mean *p*CO_2_ of approximately 460 μatm. While this is higher than current atmospheric levels it is not uncommon for the average *p*CO_2_ near shallow coral reef areas to be higher than atmospheric CO_2_ because of the combined effects of calcification from corals and respiration by reef organisms, especially during the summer months when this study was conducted (Albright et al., 2013; Page et al., 2016; Page, Courtney, Collins, De Carlo, & Andersson, 2017). Reducing the pCO_2_ of incoming water was only possible for the control treatments and consequently the lowest *p*CO_2_ levels in the 750 ± 300 μatm treatment matched the pCO_2_ of the incoming seawater (500-550 μatm). Furthermore, it was not possible to reduce *p*CO_2_ levels below 400 μatm and thus our control fluctuating treatment has a higher average *p*CO_2_ of 550 with fluctuations of +/- 150 μatm only. Mean values for seawater parameters in the experiment are presented in Supplementary Table S1.

### Extractions and RNA sequencing

At the age of ten weeks, for both species, eighteen individuals from each clutch (three fish per treatment) were randomly selected to obtain brain tissue for gene expression analysis. With three clutches for each species, the sample size was nine individual replicates per treatment for each species. With six CO_2_ treatments this resulted in a total sample of 54 individuals per species (Supplementary Table S2). Fish were taken directly from the CO_2_ treatment tanks without behavioural testing to measure the molecular response to the CO_2_ treatment rather than the behavioural and handling response. Fish in the fluctuating CO_2_ treatments were sampled at the lowest point in the pCO_2_ cycle as the as the fluctuations pass the mean level in the 750 μatm and 1000 μatm for a comparison with the stable 750 μatm and 1000 μatm treatments. All biological samples were collected at the same time point a day to avoid biases due to time of day (Figure 1).

Fish were sacrificed by severing the spinal cord and were immediately snap frozen in liquid nitrogen. Whole brain tissue was dissected out the brains under a dissection microscope. The tissue was placed into 350μl of RLT Buffer from a Qiagen All prep DNA/RNA kit. Total RNA extractions were performed (after DNA removal and extraction) according to the manufacturer’s instructions. RNA quantity was measured on a nanodrop and quality was assessed with an Agilent Bioanalyzer and samples with a RIN of 8 and above were accepted. RNA sequencing library prep was performed with Illumina TruSeq kits and sequenced on a HiSeq 4000 for paired end reads of 150bp length at Macrogen, South Korea.

### Sequence processing and differential expression analysis

Sequencing reads were processed (Supplementary Figure S2) the same way for both species with the exception of a different reference genome for each species with the relevant annotations. For *Ac. polyacanthus* the *de novo* assembled reference genome used was under BioProject PRJNA311159 and for *Am. percula* the reference genome is BioProject PRJNA436093 and metrics as well as details can be found in Lehmann et al. (2019). Raw fastq sequencing reads were trimmed with Trimmomatic (Bolger, Lohse, & Usadel, 2014) using the parameter sliding window 4:20 and removing Illumina adapter sequences. Quality-trimmed reads were then mapped against the relevant genome reference with Tophat2 v2.0.9 (Kim et al., 2013) with the ‘b2-very-sensitive’ parameter. Resulting bam files were then used to obtain raw read counts with ht-seq v0.6.1 (Anders, Pyl, & Huber, 2014) to receive transcript level counts. Differential gene expression was then evaluated with Deseq2 (Love, Huber, & Anders, 2014) package in R. Likelihood Ratio Tests (LRT) were performed for each species to evaluate the general expression patterns across all CO_2_ treatments while evaluating the influence and factoring in the family line (model= ~Family+ CO_2_ Treatment; Supplementary Figure 3a&b). Furthermore, pair-wise comparisons between CO_2_ treatments were computed by Wald tests. Transcripts with FDR corrected values of <0.05 and a minimum log2fold change of 0.3 were accepted as differentially expressed. We used Blast2Go Pro (Conesa et al., 2005) for functional enrichment tests using Fisher’s exact tests. Subsets of differentially expressed genes were therefore compared with the rest of the gene set in the transcriptome of the corresponding species. Common differentially expressed genes and functions are based on overlap in the gene name from the gene annotation in the respective genomes as both genomes were assembled and annotated the same way (R. Lehmann et al., 2019, 2021).

## Results

### Common responses across species

We measured gene expression levels across the genomes of both study species and tested for differential expression among different CO_2_ levels and treatments. When all samples and treatments are analyzed together the number of differentially expressed genes differed between the two species, with *Ac. polyacanthus* exhibiting 52 differentially expressed genes, where *Am. percula* exhibited only one differentially expressed gene (Supplementary Table S3) even though *Am. percula* displayed a wider range of fold changes (Supplementary Figure 4). This small amount of differential gene expression when including all samples and treatments shows that the fish in the different treatments react differently and do not have a common pattern of differential expression across treatments. Pairwise comparisons of treatments revealed a greater number of differentially expressed genes (Figure 2). There was overlap among the two species in the transcriptional response to elevated stable CO_2_ in pairwise comparisons among treatments. Comparing the reaction to stable high CO_2_ treatments (450 vs 750 or 1,000 μatm) 20% of the genes differentially expressed (DEGs) in *Am. percula* were also differentially expressed in *Ac. polyacanthus* (Supplementary Table S4) indicating some overlap in differential expression between these species. The common response to elevated stable CO_2_ in both species was downregulation of immediate early genes (IEG; Pérez-Cadahía, Drobic, & Davie, 2011) such as Fos-Proto-Onco gene (FOS) or Jun-Proto-Onco gene (JUN; Supplementary Table S5). Other common genes differentially expressed in both species are involved in glycolysis such as lactate dehydrogenase (ldha) and midnolin (midn). These similarities in differential expression between the species occurred despite differences in the collection locations of adults and the duration of time that breeding pairs of each species had spent in captivity.

**Fig. 2:**
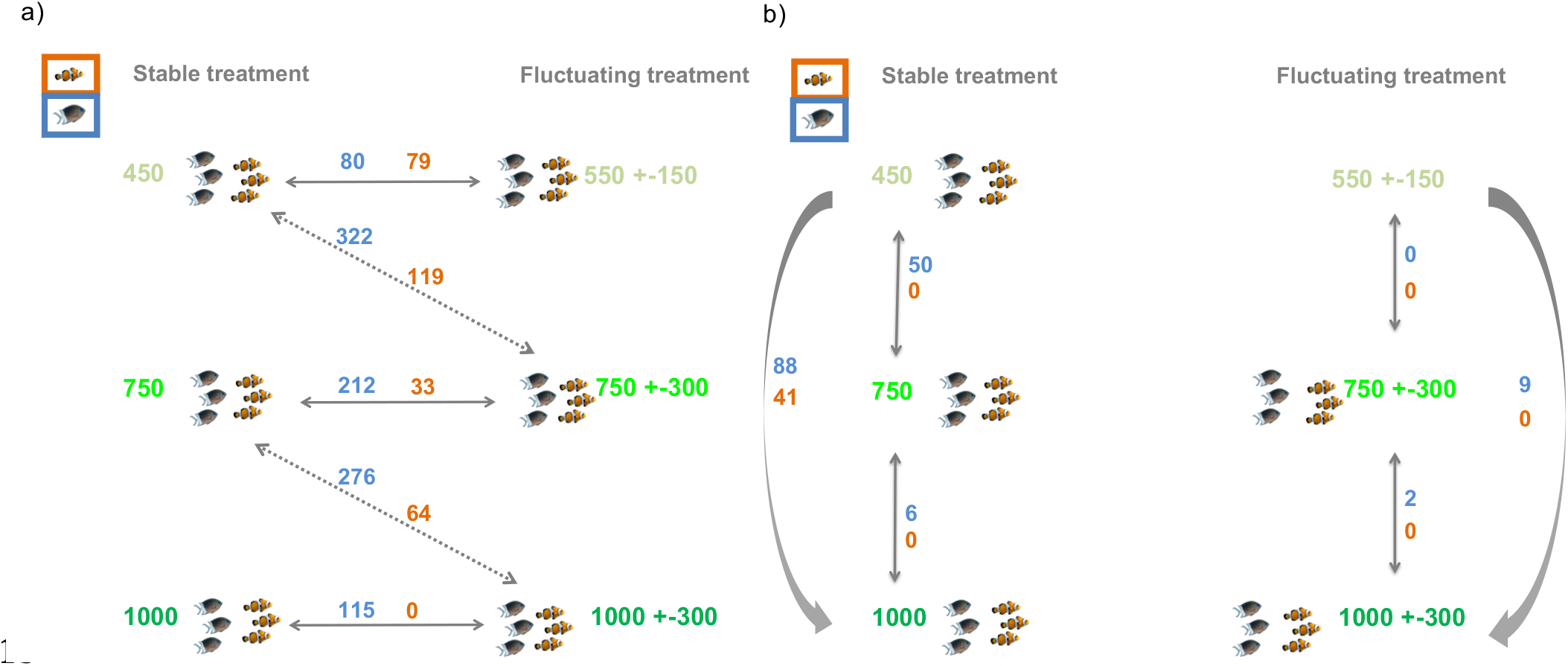
Number of genes that are differentially expressed between different treatments in *Acanthochromis polyacanthus* (blue) and *Amphiprion percula* (orange) comparing a) across stable and fluctuating treatments. Solid arrows represent comparisons with similar average pCO_2_ levels but different levels at the time of collection whereas dashed arrows represent comparisons between similar levels of environmental pCO_2_ at the time of collection. b) number of differentially expressed genes within stable or fluctuating treatments but with different levels of average pCO_2_.

### Gene expression in stable versus fluctuating CO_2_ treatments

Transcriptional regulation differed for fish exposed to stable and fluctuating CO_2_ treatments for both *Ac. polyacanthus* and *Am. percula*. The direct comparison between stable and fluctuating CO_2_ treatments resulted in the largest difference in gene expression for both species (Fig. 2a). We were able to look at gene expression differences for two types of comparisons. Firstly, we compare stable and fluctuating treatments with the same average levels of CO_2_ per treatment (Fig. 2a, solid arrows). These comparisons, regardless of the average CO_2_ being at 450, 750 or 1000μatm, resulted in similar functional changes in the brain of our two fish species. Changes between fluctuating and stable CO_2_ treatments resulted in differential regulation of the response to steroid hormones and the circadian rhythm. *Ac. polyacanthus* also had enriched functions involved in oxidation-reduction (Supplementary Table S6), whereas *Am. percula* differentially expressed genes involved in gluconeogenesis (Supplementary Table S7). However, for these treatment comparisons, while having the same average CO_2_ levels across the day, at the moment of collection the environmental levels of CO_2_ differed (Fig. 1).

A second set of comparisons were made for fish that were at similar levels of environmental CO_2_ at the time of sample collection, while the average CO_2_ level across the day differed (Fig. 2a, dashed arrow). Despite the same level of CO_2_ at time of collection, these comparisons resulted in the largest gene expression differences. For the comparison between control stable treatment (450 μatm) and the fluctuating treatment of 750+-300 μatm there were 322 genes differentially expressed for *Ac. polyacanthus* and 119 for *Am. Percula* (Fig. 2a). Similarly, when comparing 750 μatm to 1,000+-300 μatm, which have similar CO_2_ levels at the time of sample collection, 276 and 64 genes for *Ac. polyacanthus* and *Am. percula* were differentially expressed. Regardless of the level of CO_2_ at time of collection, when comparing between stable and fluctuating treatments some genes were always differentially expressed. There were sixteen such genes for *Am. percula* (Supplementary Table S8) and 107 for *Ac. polyacanthus* (Supplementary Table S9), six of which are common among the two species (PYRG1, HLF, BHLHE40, NFIL3, NPAS2/4, NR1D2/4). The functions underlying the common genes that differ between stable and fluctuating environment are involved in processes of circadian rhythm as well as steroid hormone signalling pathway in both species (Fig. 3).

**Fig. 3:**
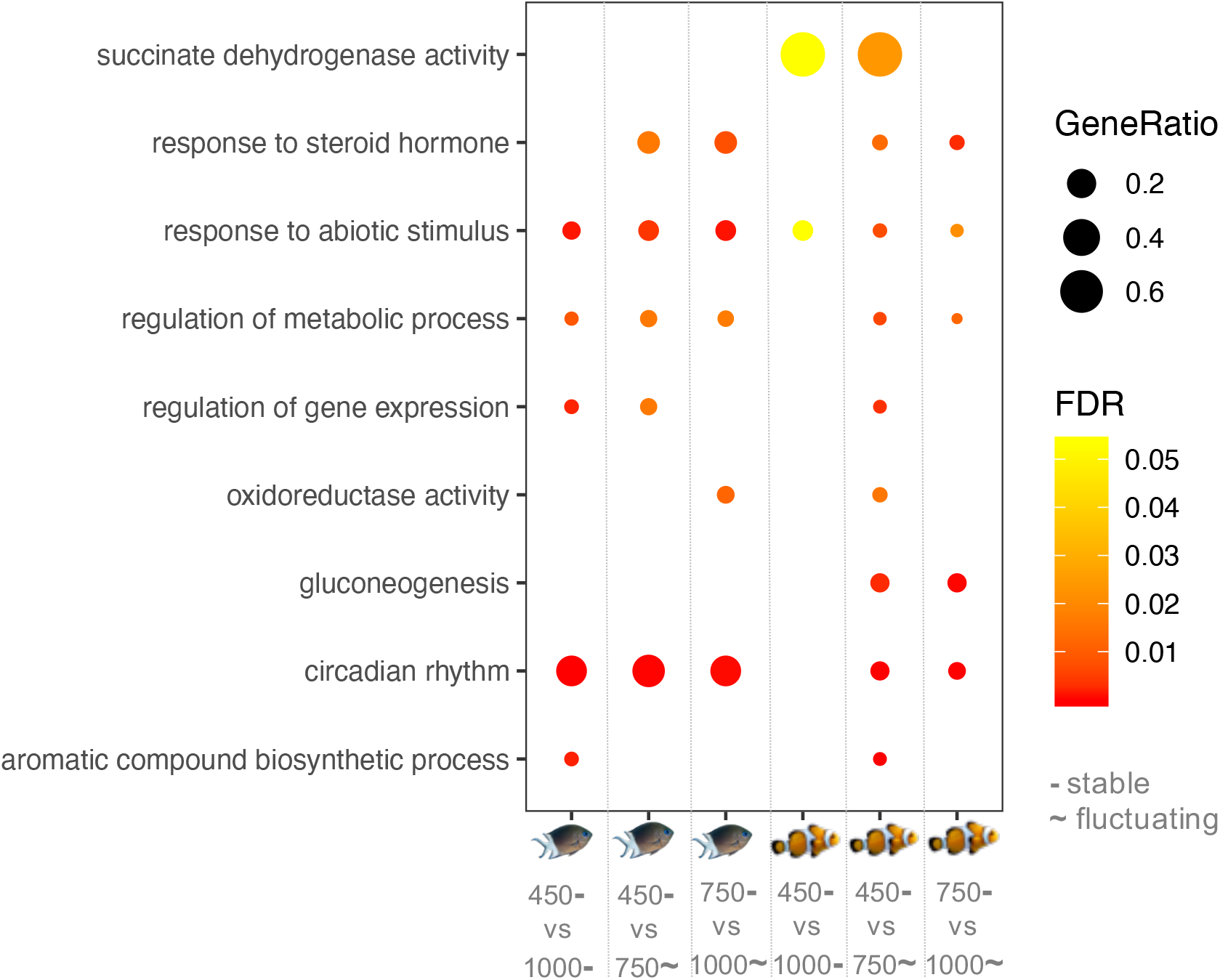
Enriched functions related to differentially expressed genes in *Acanthochromis polyacanthus* and *Amphiprion percula* for three different CO_2_ treatment comparisons per species. FDR represents statistical significance level after FDR adjustment of the enrichment with red indicating higher significance. Gene ratio is the number of genes represented in the differentially expressed genes in comparison to the rest of the transcriptome.

### Gene expression among CO_2_ treatment levels

Despite some common differentially expressed genes, the magnitude of response differed between the two species. *Ac. polyacanthus* had a larger molecular reaction with more differentially expressed genes across all stable CO_2_ treatment comparisons than *Am. percula* (Fig. 2b), with a total of 88 DEGs for *Ac. polyacanthus* and 41 DEGs for *Am. percula*. In fact, *Am. percula* exhibited no change in transcription when comparing 450 μatm to 750 μatm or 750 μatm to 1,000 μatm in both stable and fluctuating CO_2_ treatments. Only the comparison between stable 450μatm and 1,000 μatm resulted in 41 DEGs for *Am. percula*. For the stable elevated CO_2_ treatment, we found no enriched functions for *Am. percula* whereas *Ac. polyacanthus* displayed enrichments in several functions including the circadian rhythm and RNA metabolic processes (Fig. 3 & Supplementary Table S10).

For the species with more gene expression changes, *Ac. polyacanthus*, an increase of CO_2_ in a stable treatment resulted in a higher number of differentially expressed genes between stable 450 to stable 750 μatm (Fig. 2b; 50 genes) than from stable 750 to stable 1,000 μatm (6 genes). The main gene expression pattern, found in 27 genes, was either an increase or decrease in expression from stable 450 to 750 μatm, followed by a levelled expression for stable 750 to 1,000 μatm (Supplementary Table S11). These genes have a variety of functions including gluconeogenesis, neuronal development and cellular stress response. In particular several immediate early genes, such as FOS, EGR1 and JUN, were expressed at lower levels in the elevated CO_2_ treatments, but also the Neuronal PAS domain protein 4 (NPAS4), particularly involved in homeostatic brain plasticity (Lin et al., 2008).

Comparing among pCO_2_ treatment levels we found that gene expression differences were more pronounced for fish from different levels of stable CO_2_ conditions than from different CO_2_ levels of fluctuating treatments, especially for *Ac. polyacanthus*. Between the stable elevated CO_2_ treatment of 1,000 μatm and the stable control (450 μatm) condition there were 88 and 41 genes differentially expressed for *Ac. polyacanthus* and *A. percula* respectively (Fig. 2b; Supplementary Table S12 & S13). For the same average CO_2_ treatments levels with diel fluctuations, only nine genes and zero genes were differentially expressed for the two species respectively (Supplementary Table S14). For *Ac. polyacanthus* one gene (BMAL1) was differentially expressed when comparing low (450/550 μatm) versus elevated (1000 μatm) CO_2_ in both the stable and fluctuating treatments. BMAL1 is one of the main circadian rhythm activators representing a response to elevated CO_2_ regardless of stable or fluctuating treatment. The reaction for *Ac. polyacanthus* among the stable treatments (88 genes) was enriched in functions such as the circadian rhythm, response to abiotic stimulus, and aromatic compound metabolism (Supplementary Table S6). The genes reacting to the difference in CO_2_ level in fluctuating treatments (nine genes) were involved in the circadian rhythm (BMAL1, NFIL3); neuronal morphology (CRLF3, NYAP2); immunity (NOD1); neural activity, cell signalling and glutamergic transmission (PDE7B, SLC7A10) and iron ion binding (CP2J2, LONRF3). Even though the magnitude in transcriptional response as well as the actual genes that are differentially expressed differ for the low to high CO_2_ comparisons within the stable and fluctuating treatment comparison, many of these genes have similar functions (Fig. 3).

### Genes in the Circadian Rhythm Pathway influenced by CO_2_ treatment

The circadian rhythm (CR) was one common mechanism affected by different CO_2_ treatments due to changes in gene expression of the pathway’s activator and repressor genes in both studied species. All genes in the pathway change expression levels with the increase in stable CO_2_ levels, whereas fluctuating treatments showed a more drastic change in gene transcription with downregulation of CR repressor genes and upregulation of CR activator genes in the fluctuating CO_2_ treatments compared with the stable CO_2_ treatments (Fig. 4). At the time of collection, which coincides with the start of the diel low peak of CO_2_ in the fluctuating treatments, the CR activators such as CLOCK and BMAL were lowly expressed for the stable 450 μatm (Control) treatment in both species (Fig. 4 a & b), as expected at this time of day. In contrast, these central activator genes to the CR had higher levels of expression in the stable elevated CO_2_ treatments with elevated expression at 1,000 μatm (Fig. 4). CR repressors, such as CRY or PER exhibited the opposing pattern, with greater expression at 450 μatm (as to be expected this time of day) and lower levels in the stable elevated CO_2_ treatment. Hence, the expression of the whole CR is affected by elevated CO_2_. However, when CO_2_ fluctuations are introduced the expression changes even more, as the magnitude of change in expression of activator and repressor genes increases markedly in the elevated CO_2_ treatments with diel fluctuations. Even comparing stable 450 μatm with fluctuating 550 μatm there was an increase in activator expression or a decrease in repressor expression (Fig. 4). The actual environmental CO_2_ levels were similar at the time of collection for the fluctuating 750 and stable 450 μatm as well as between fluctuating 1000 and the stable 750 μatm, however there were large differences in gene expression between those treatments for the CR genes. This shows that it is not the time of day nor the momentary enviromental CO_2_ levels that are driving the CR gene expression patterns as all were the same between treatments.

**Fig. 4:**
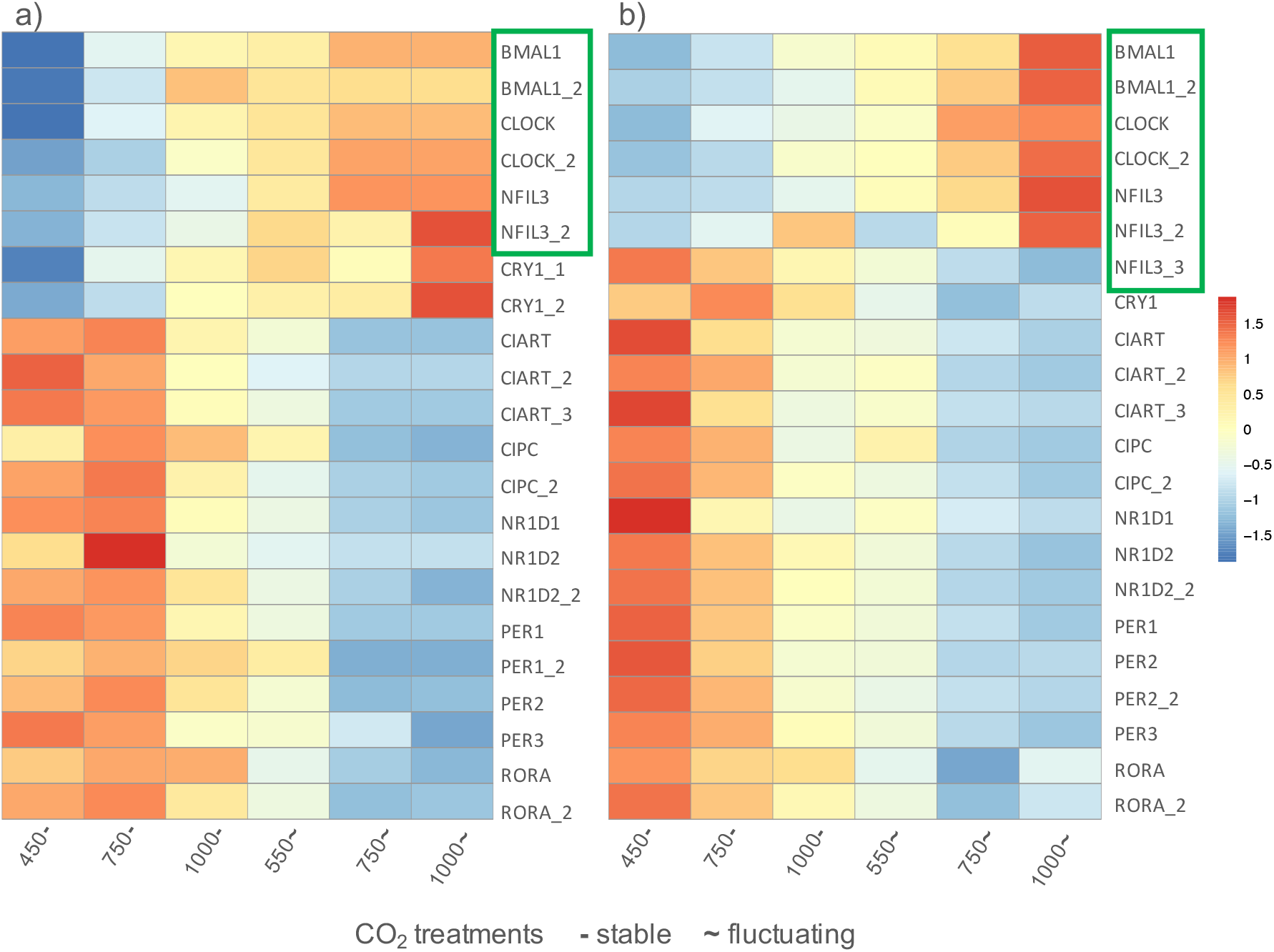
Gene expression signal of Circadian rhythm (CR) genes across the six different CO_2_ treatments for a) *Acanthochromis polyacanthus* and b) *Amphirion percula* based on average normalized read counts per treatment. Differential expression for each pairwise comparison can be found in Supplementary Table S15 & 16). Green boxes indicates CR ‘activator genes’ whereas other genes are ‘repressor genes’ of the CR. Heatmap is represented on a z-score with lower expression in blue and elevated expression in red.

## Discussion

The molecular adjustments made in brains of fish when exposed to elevated CO_2_ levels mimicking future ocean acidification show some, albeit limited, similarities in genes as well as functions across two coral reef fish species. We found a set of genes known as immediate early genes (IEGs) differentially expressed in the brain of both species when exposed to stable elevated CO_2_. In a previous study some of these genes were also found differentially expressed in *Ac. polyacanthus* brains after an acute 4day elevated CO_2_ treatment, but mostly for offspring of behaviourally sensitive individuals (Schunter et al., 2018). Expression of a number of immediate early gene is commonly used to understand neuronal activity for certain behaviours in vertebrate species (Guzowski et al., 2005). Increased expression of these genes is an indicator for neural activity especially tied to a specific behaviour or external stimulus with a variety of downstream functions (Pérez-Cadahía et al., 2011). In other fish species, increased expression was shown as a response to neurotoxin from algal blooms (Salierno et al., 2006), exposure to nickel (Topal et al., 2015), a sign of parental care (O’Connell, Matthews, & Hofmann, 2012) and with aggression or feeding (Wai, Lorke, Webb, & Yew, 2006). We found that expression of early immediate genes decreased in elevated CO_2_ treatments compared with control CO_2_. While expression of immediate early genes, such as C-FOS, is often detected in certain areas of the brain only, we examined whole brain gene expression and still found generally lower levels of expression in the whole brain tissue of fish exposed to elevated CO_2_. A decrease in C-FOS expression has received little attention, as mostly increased expression is used to localize the response to a stimulus in the brain. However, an expression decrease has been associated with lower aggression levels as well as social subordinance in another *Amphiprion* species (Yaeger et al., 2014). As IEGs are used as indicators of brain activity, it is possible that the decreased expression of these genes in all elevated CO_2_ treatments, for both species, suggests a lower level of brain activity related to this environmental stimulus. In fact, a decrease in activity in various essential behaviours in fish, such as locomotion and feeding behaviour have been linked to a decrease in brain activity when exposed to toxins (Berntssen, Aatland, & Handy, 2003; Rao, Begum, Pallela, Usman, & Rao, 2005). As previously reported in Jarrold et al. (2017) the fish in thus experiment exhibit altered lateralization (turning preference) and impaired reaction to chemical alarm cues when exposed to elevated CO_2_. The change in behaviour was strongest with an increase in stable CO_2_, and while whole brain gene expression is not necessarily linked to a particular behaviour, the lower expression of brain activity genes could lead to behavioural changes.

*Ac. polyacanthus* (spiny chromis damselfish) exhibited more transcriptional changes than *Am. percula* in response to elevated CO_2_ in the brain, even if the transcriptional response found here is smaller than previously reported for acute exposures at the 750 level μatm (C. Schunter et al., 2018). Compared with other coral reef fishes, the spiny chromis also exhibits a larger transcriptional change to thermal stress (Bernal et al., 2020) and our results suggest that this sensitivity of molecular responses to environmental change applies similarly to changes in pCO_2_. Here we used two different CO_2_ treatment levels that are relevant to different ocean acidification scenarios. Our results show that the spiny damselfish is sensitive to a relatively small increase in pCO_2_, from 450 to 750 μatm, and that most of the molecular changes occur by 750 μatm, not in the additional increase to 1000 μatm. Therefore, the molecular responses of these fish are highly sensitive to even relatively small changes in pCO_2_. It was previously thought that much higher CO_2_ levels are needed before changes in bicarbonate and chloride to compensate for a pH change would occur. However, plasma CO_2_ has been shown to change at levels of 1000 μatm and increased bicarbonate levels in the plasma and brain have been shown at 1900 μatm (Esbaugh et al., 2016; Rachael M. Heuer et al., 2019). No such measurements were obtained at 750 μatm, but this is consistent with other studies showing that active acid-base regulation is engaged in some fishes at 750 μatm (Tresguerres, Milsom, & Perry, 2019). Furthermore, behavioural impairment occur in a number of reef fishes species at this CO_2_ level (Devine, Munday, & Jones, 2012; Ferrari et al., 2011; Munday et al., 2010a; Welch et al., 2014). For instance, a coral reef cleaner wrasse exhibited variable effects of CO_2_ on learning behaviour at 750 μatm but impairment occurred in all individuals at 950 μatm (Paula, Baptista, et al., 2019), also suggesting 750 μatm to be a tipping point for behavioural impairment in reef fishes. The increase in CO_2_ level from 450 to 750 initiated transcriptional responses in immediate early genes as well as genes involved in brain metabolism and neuronal development, but chloride or bicarbonate transporters did not change in expression. It is possible that other molecular processes such as post-transcriptional or translational play a role here or that transcription of such genes might occur at the beginning of the exposure to the environmental change, but after eight weeks of exposure the transcriptional program is focussed on neuronal activity and brain metabolism such as seen in previous studies on the transcript as well as the protein level (Araújo et al., 2018; Schunter et al., 2016; Tsang, Welch, Munday, Ravasi, & Schunter, 2020).

While the overall increase in average pCO_2_ is important, we found that the stability of the environmental conditions a fish experiences seems to matter more as the comparison between fish living in a stable and a fluctuating environment resulted in the largest differences to the transcriptional brain program. This indicates that the response to a change in environmental CO_2_ conditions is different for organisms living in a fluctuating environment than in stable environments. Acute exposures to more extreme environmental changes (such as high temperature or low pH) could increase individual performance in fluctuating environments in comparison to stable environments (Niehaus, Angilletta, Sears, Franklin, & Wilson, 2012). On the physiological level, organisms need to maintain their internal physiological state with a change in environmental conditions, and in the case of ocean acidification adjust their acid-base balance to avoid acidosis (Brauner & Baker, 2009). In a diel fluctuating environment this would result in a repeated adjustment to assure physiological functioning. We find the largest expression differences between stable and fluctuating environments, and these transcriptional differences disappear under different fluctuating CO_2_ levels which has similarly been seen for the behaviours measured for our two study species when comparing stable and fluctuating CO_2_ treatments (Jarrold et al., 2017). On the behavioural level there are no significant differences in lateralization among the different fluctuating treatment showing that temporary overlap of CO_2_ levels is more influential on all levels of response than average CO_2_ differences (see also Jarrold & Munday, 2019). Such a pattern can also be seen on the physiological level, where aerobic scope and oxygen uptake rates differ between two stable CO_2_ treatments, however when diel fluctuations are added, both metabolic measures are not different to control (Laubenstein et al., 2020). Previous studies on *Acanthochromis polyacanthus* (Schunter et al., 2018, 2016) suggest that transcriptional regulation might be involved in maintaining homeostasis in the fish in a stable environment while causing downstream behavioural impairments. In fluctuating environments, however regardless of CO_2_ level, transcription does not differ, and crucial behaviours are mostly maintained. Hence, for coral reef fishes CO_2_ fluctuations might allow the fish to anticipate environmental changes in pH and cope with the changes rather than adjusting the transcriptional and physiological response underlying the acid-base control.

Adaptative responses to varying environments can arise from feed-forward approaches preparing the organism to upcoming predicted changes (Bernhardt et al., 2021). One important feed-forward mechanism is adjusting the Circadian Rhythm. Both study species differentially transcribed genes involved in regulating the circadian clock confirming its fundamental importance in these processes. There was an overall downregulation of circadian rhythm repressor genes and upregulation of circadian rhythm activator genes in the fluctuating CO_2_ treatments compared with the stable CO_2_ treatments. A previous study on *Ac. polyacanthus* found that the circadian clock was differentially regulated in offspring of behaviourally tolerant parents to elevated CO_2_ (Schunter et al., 2016) suggesting a more flexible approach to acid-base regulation. It seems that in fluctuating environments the circadian clock shifts downstream physiological processes, such as growth, by many hours (Nozue et al., 2007). This process likely applies for reef fishes as the circadian cyclical genes, represented by activators and repressors, were shifted. Such a circadian phase shift was also encountered for the olive flounder with different levels of elevated environmental CO_2_ (Choi, Lee, Song, & Park, 2021). Furthermore, a regulatory connection between the Circadian Rhythm and extracellular pH was found in the goldfish retina concluding that oscillations/fluctuations control neuronal activity and pH (Dmitriev & Mangel, 2000). Changes to neuronal activity in addition to the CR expression changes, were also observed in the brain of the two coral reef fishes hinting to a relationship between changes in environmental CO_2_ and neuronal activity with CR oscillations. However, this requires future studies across different times of the circadian day to verify the connection between environmental pH changes, brain activity and the Circadian Rhythm. In this study we found that diel fluctuations in CO_2_ in coral reef systems allow coral reef fishes to adapt to constant changes in the environmental pH and exhibit flexibility in adjustments to such pH changes most likely driven by the circadian rhythm regulator in the brain. Adaptations to diel fluctuations might therefore alleviate negative effects on coral reef fish species even with elevated CO_2_ conditions expected due to ocean acidification.

## Acknowledgments

This study was supported by the Office of Competitive Research Funds OSR-2015-CRG4-2541 from the King Abdullah University of Science and Technology (T.R., P.L.M., C.S.), the Australian Research Council (ARC) and the ARC Centre of Excellence for Coral Reef Studies (P.L.M). This project was completed under approval of the James Cook University animal ethics committee (permit: A2210) and according to the University’s animal ethics guidelines. We thank the Marine and Aquaculture Research Facilities Unit (JCU) and the Integrative Systems Biology Laboratory (KAUST). A special thanks goes to Damien Lightfoot for extracting RNA from *Am. percula* brains and Robert Lehmann for aiding with the genome mapping of *Am. percula* sequences.

## Author contributions

M.J. and P.L.M designed and managed the fish rearing experiments. C.S. prepared the samples for RNA sequencing, analysed the transcriptome data and wrote the first draft. All authors edited and approved the final manuscript.

## Additional information

RNA-seq transcriptome raw sequences have been deposited in NCBI under BioProject PRJNA657978 for *Amphiprion percula* and PRJNA657977 for *Acanthochromis polyacanthus*.

