## Supplementary Figures for "Diel CO_2_ fluctuations alter the molecular response of coral reef fishes to ocean acidification conditions"

**Diel CO_2_ fluctuations affect the brain transcriptome of coral reef fishes under ocean acidification conditions**

SUPPLEMENTARY FIGURES:

Supplementary Figure 1:

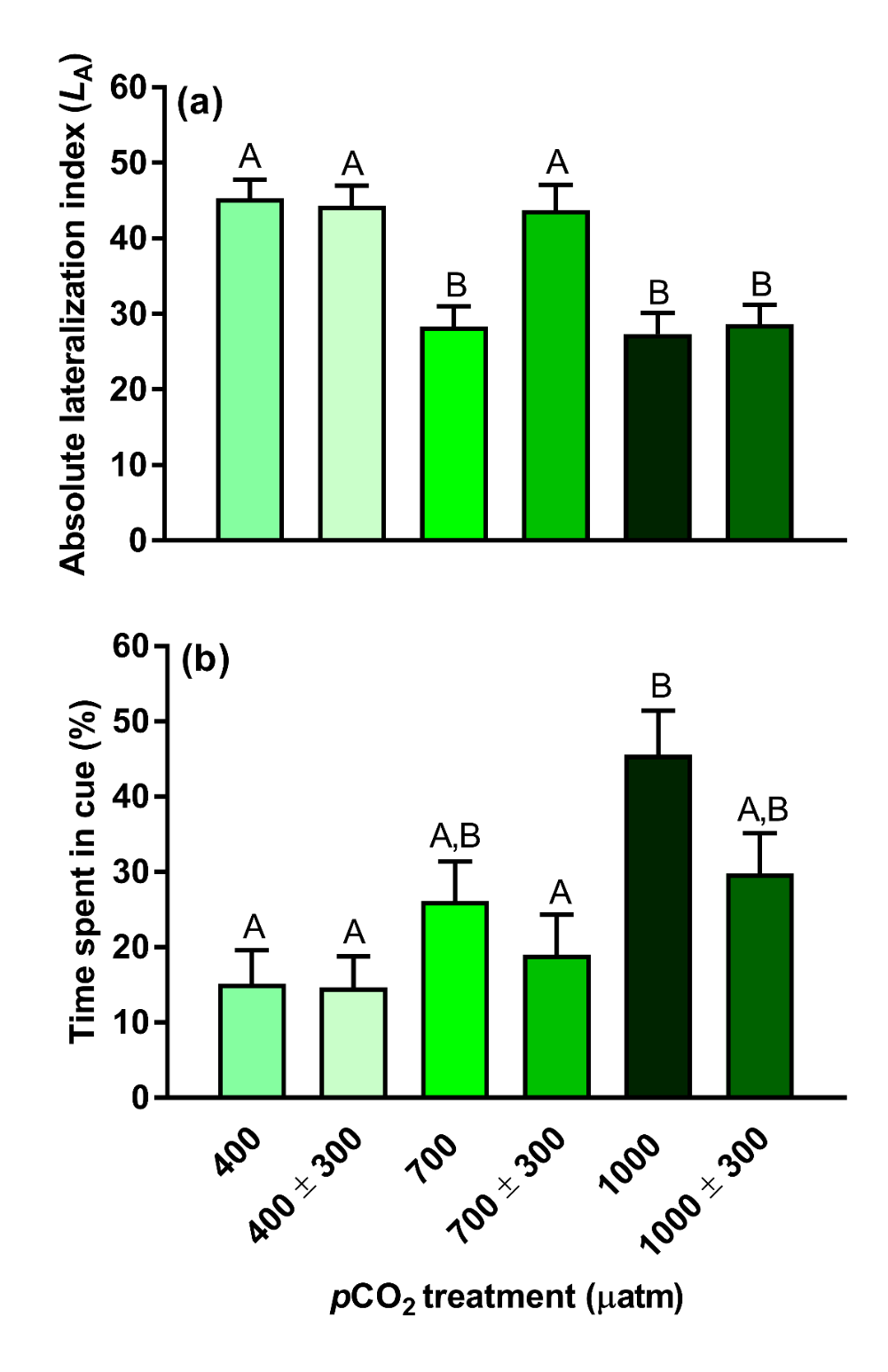

Figure S1: The effect of pCO_2_ treatment on (a) absolute lateralization of juvenile *Acanthochromis polyacanthus* (n = 60 per treatment) and (b) response to predator cue of juvenile *Amphiprion percula* (n = 24 per treatment) in experiment two. Different letters represent significant differences between treatments (Tukey, P < 0.05). Bars are mean values ± SE.

Supplementary Figure 2:

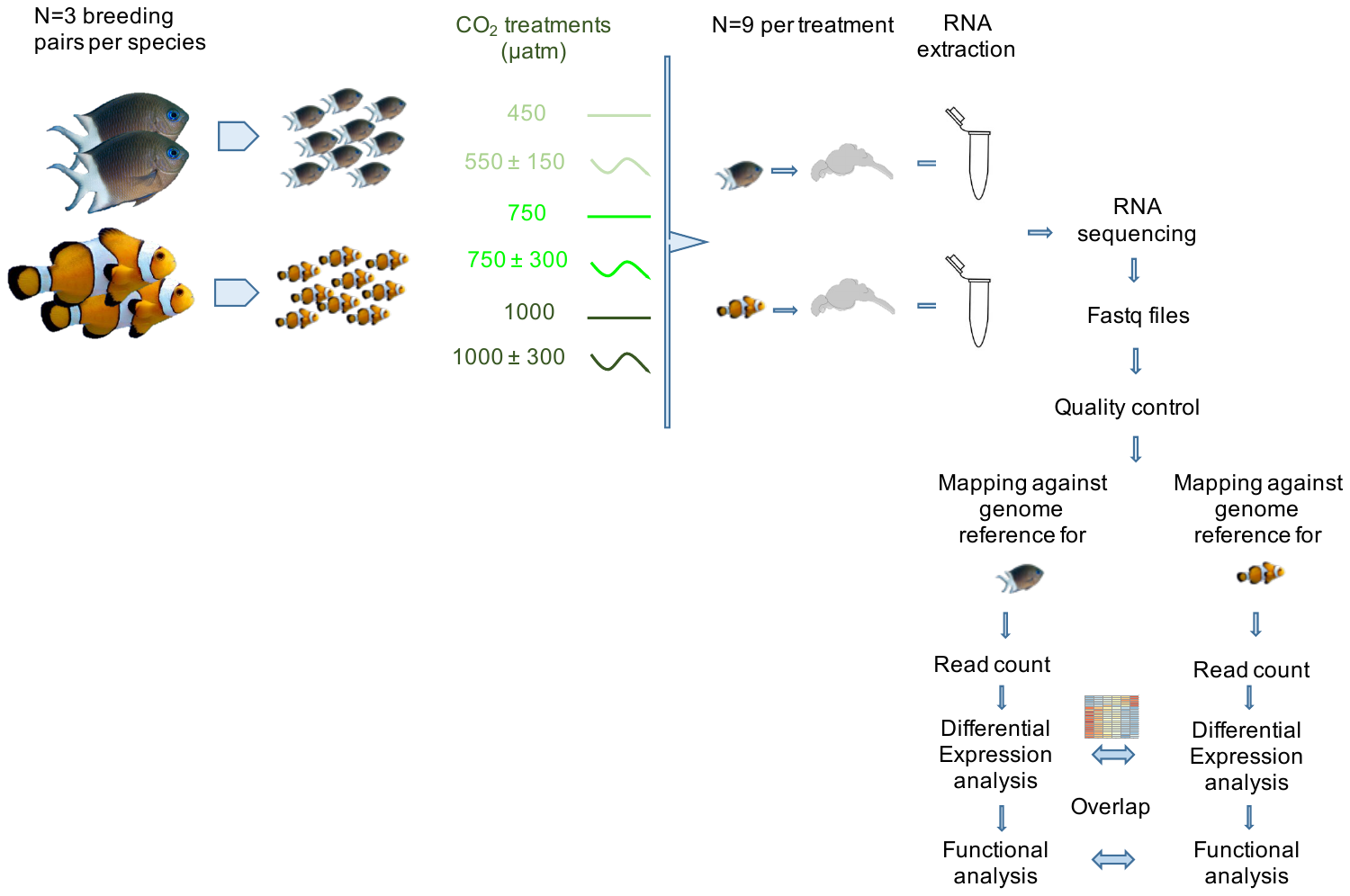

Figure S2: Experimental design including bioinformatic pipeline. Juveniles of the two species were exposed to six different CO_2_ treatments (3 stable and 3 fluctuating), sacrificed and brains dissected. Brain RNA was sequenced, and high-throughput sequencing files were processed for quality and mapped against the respective genome reference. Differential expression by comparing all pair-wise treatment groups and functional analyses was performed for each species separately and overlap was determined by differentially expressed genes and enriched functional categories.
Supplementary Figure 3:

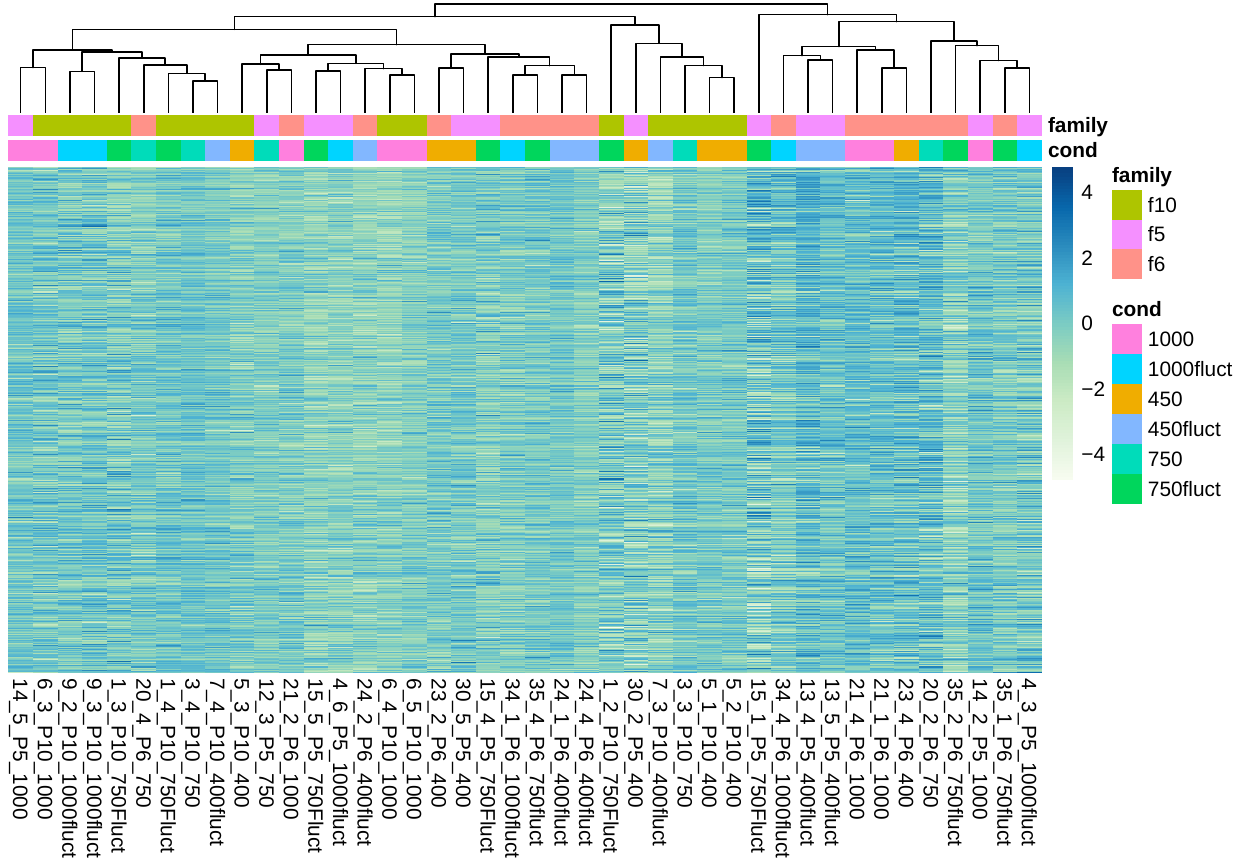

Supplementary Figure S3: Heatmap of individuals of *Acanthocromis polyanthus* clustering indicating CO_2_ treatments and family line.

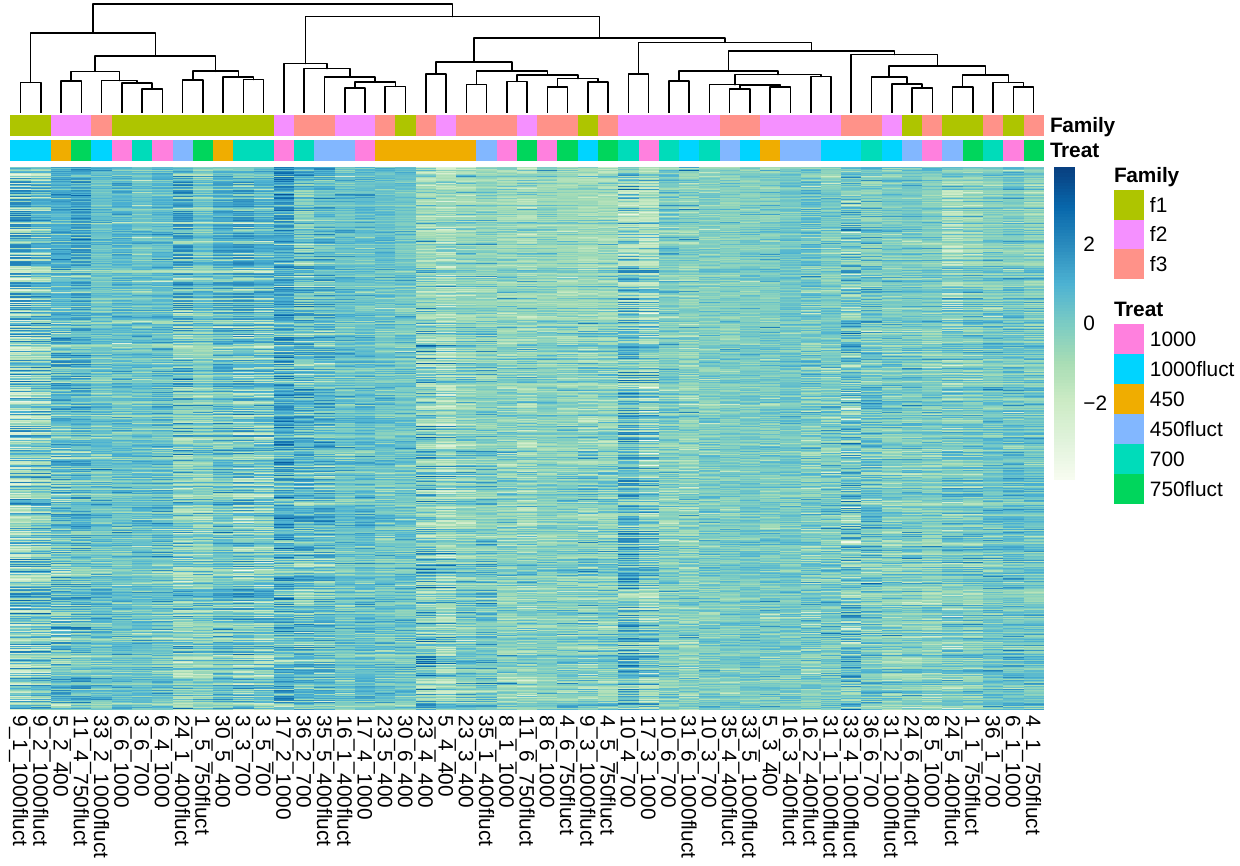

Supplementary Figure S3b: Heatmap of individuals of *Amphiprion percula* clustering indicating CO_2_ treatments and family line.

Supplementary Figure 4:

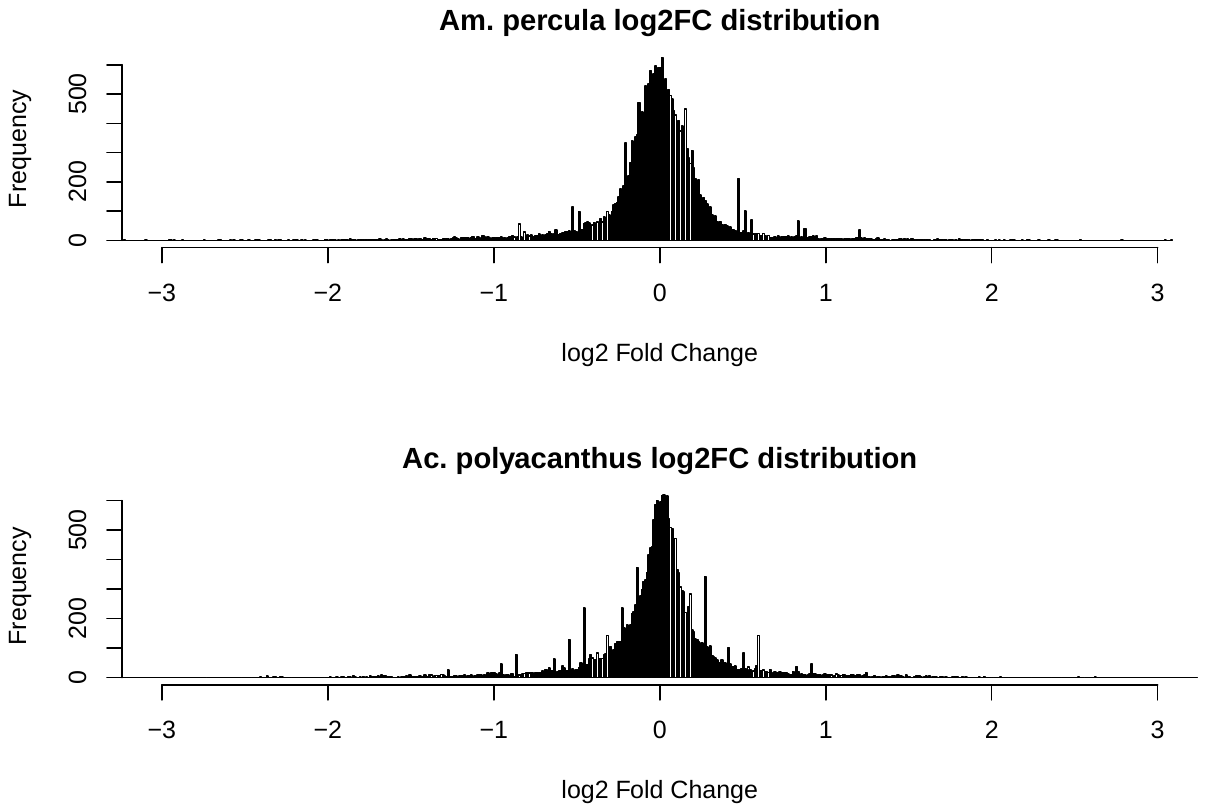

Supplementary Figure S4: Distribution of Log 2-fold changes across the whole transcriptome for *Amphiprion percula* (top) and *Acanthochromis polyacanthus*
